# Single-Cell Network Analysis Identifies CLEC4E as a Key Mediator of Proinflammatory mDC Responses in Influenza Infection

**DOI:** 10.1101/2025.08.21.671587

**Authors:** Subin Cho, Gabriel Laghlali, Arturo Marin, Adolfo García-Sastre, Gagandeep Singh, Michael Schotsaert, Won-Min Song, Christian V. Forst

## Abstract

The severity of influenza is often driven by an excessive host immune response rather than the virus itself, yet the key molecular drivers within specific immune cells remain poorly understood. While recent single-cell RNA sequencing studies have successfully identified immune populations involved, they have largely not identified the upstream drivers modulating their pro-inflammatory functions. Here we employed an integrated single-cell co-expression network to address this gap. Our analysis identified myeloid dendritic cells (mDCs) as central to pro-inflammatory response during infection. Through a multi-layered key driver analysis, we pinpointed C-type lectin, *CLEC4E* as a top candidate modulating this pathological inflammatory response. The role of *CLEC4E* was confirmed in an independent single-cell dataset from influenza-infected patients and further validated *in vivo*. Pharmacological inhibition of *CLEC4E* in a murine influenza model significantly reduced disease severity and lower viral titers in the lungs. This study not only clarifies that *CLEC4E* overexpression in mDCs contributes to pro-inflammatory signaling pathways influencing influenza severity but also shows the power of single-cell network approaches to uncover novel and robust therapeutic targets hidden within complex immune responses.

## Introduction

Influenza virus infection continues to pose a significant global public health challenge^1–3^. While the host immune response is necessary for facilitating recovery, the severity of this disease is often dictated more by the host’s excessive immune response than by the virus’s intrinsic pathogenicity^4^. This dysregulated inflammatory response can precipitate severe immunopathological outcomes, such as acute lung injury^4–6^ which is driven by the complex interplay of innate immune cells during the early phases of infection^7–9^. Therefore, understanding and controlling the host-side factors that trigger excessive harmful inflammation is a promising avenue for developing effective therapies against severe influenza.

Elucidating the contributions of specific cell subtypes within complex immune reactions has been challenging with traditional bulk analysis methods. The advent of single-cell RNA sequencing (scRNA-seq) technologies has enabled the analysis of cellular heterogeneity and cell-specific responses during infection with unprecedented resolution. Indeed, scRNA-seq studies of respiratory viral infections like influenza and COVID-19 have consistently highlighted myeloid cells, particularly monocytes and myeloid dendritic cells (mDCs) as major producers of inflammatory cytokines and chemokines, playing pivotal roles in immunopathogenesis^10–12^. For instance, a recent study by Lee *et al.*^13^ identified inflammatory signatures driven by type I interferon and TNF/IL-1β specifically within classical monocytes during infections from influenza and COVID-19 patients. Similarly, Zhang *et al.*^14^ generated a comprehensive single-cell atlas of peripheral immune response in influenza A virus infected patients, further clarifying the roles of diverse immune populations.

While these studies have been instrumental in identifying the key cellular players, their analyses largely conclude with the identification of differentially expressed genes (DEGs) and their functions. This approach, though informative, highlights individual gene changes but fails to capture the coordinated regulatory logic that governs cellular behavior. Consequently, the upstream key molecular regulators which modulate the inflammatory functions of these myeloid populations remain largely unidentified. Identifying such regulators is essential for developing targeted therapeutic strategies that can control inflammatory responses. Co-expression network analysis offers a path beyond simple DEG lists by modeling the intricate network of regulatory relationships. This systems-level approach is essential for identifying functional modules and pinpointing the key driver genes that act as central hubs in the host immune response.

Therefore, to move beyond the limitations of DEG centric analyses and discover the key upstream drivers, this study employed an integrative strategy that applies co-expression network analysis to scRNA-seq data from influenza patients. First, we identified cell-type specific gene responses to viral infection and constructed co-expression networks to find out which cell types affect the most during infection. Next, using an integrated multi-layer analysis, we systematically identified key driver genes which regulate the primary immune responses (*Figure 1*).

**Figure 1.**
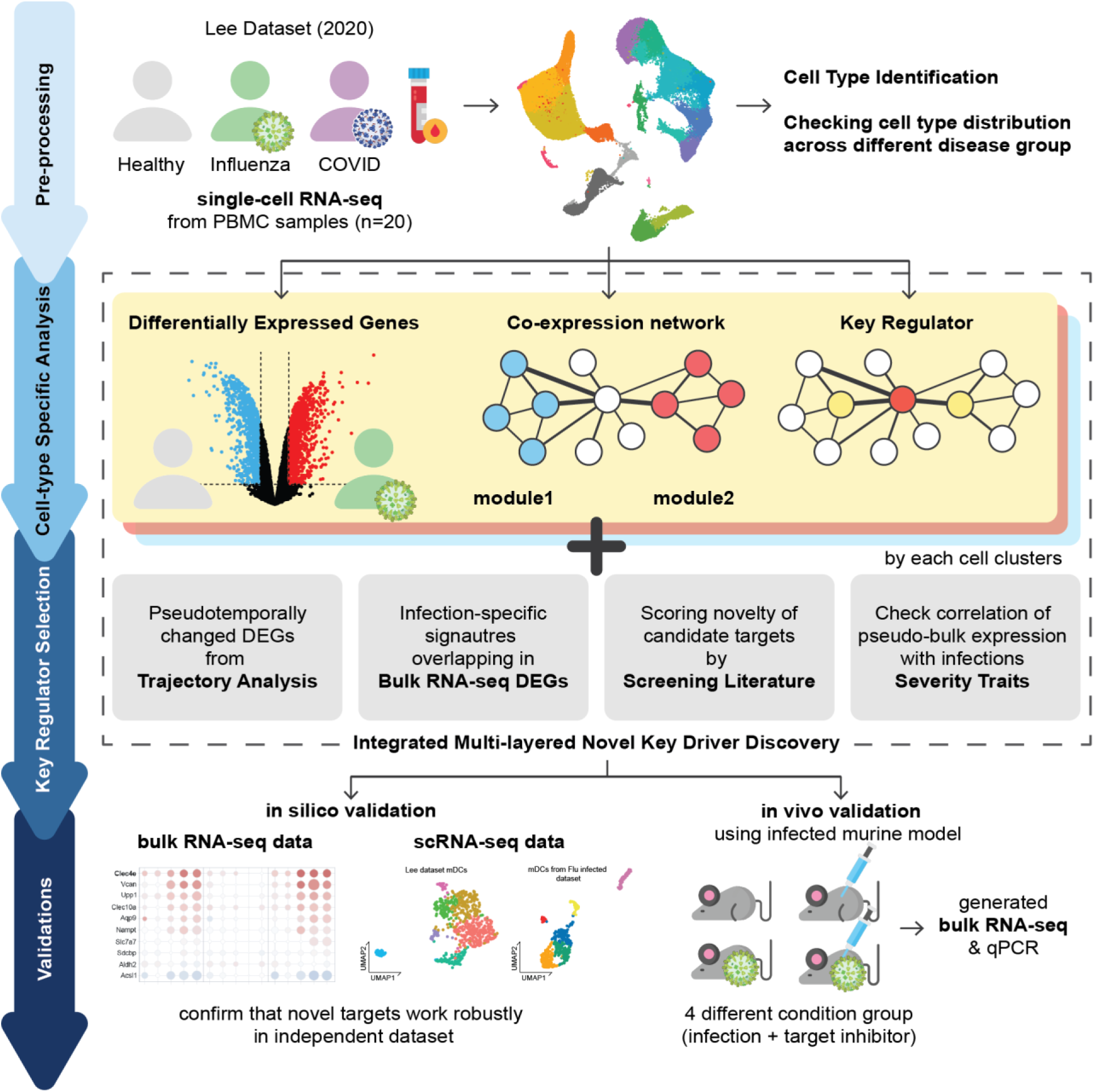
Study Overview illustrates the multi-step pipeline used in this study. The workflow starts with the pre-processing of scRNA-seq data from peripheral blood mononuclear cells(PBMCs) of healthy controls, influenza patients and COVID-19 patients. And then it’s followed by cell-type specific analysis and key regulator selection stage. This approach combined analyses of differentially expressed genes(DEGs), co-expression networks, key regulators, pseudotemporal trajectories, correlation check of infection traits with information from bulk RNA-seq and literature screening to score and prioritize candidate target genes. Finally, top candidate target were subjected to validation, including *in silico* confirmation using independent datasets and functional validation *in vivo* using an infected murine model with pharmacological inhibitor.

This approach led us to identify the C-type lectin. *CLEC4E* is a top-ranked novel regulator driving pro-inflammatory responses specifically in mDCs. We subsequently validated our findings through both *in silico* analysis of independent datasets and *in vivo* experiments. This study not only suggests a novel role for *CLEC4E* in influenza immunopathology but also demonstrates the power of single-cell network analysis for uncovering novel therapeutic targets.

## Results

To characterize the host immune response to infectious disease at single-cell resolution, we performed an in-depth analysis of single-cell transcriptomes of 62,290 cells from peripheral blood mononuclear cells (PBMCs). The dataset comprised of samples from influenza infected patients (n=5), SARS-CoV-2 infected patients with varying range of severity (n=11), and healthy controls (n=4) from a public study by Lee et al. 2020 ^13^ (hereafter referred to as the Lee dataset). Following rigorous quality control, we applied an unsupervised clustering and integration algorithm which partitioned cells into 19 distinct clusters based on their transcriptional profiles (**Figure 2 a, b**).

**Figure 2.**
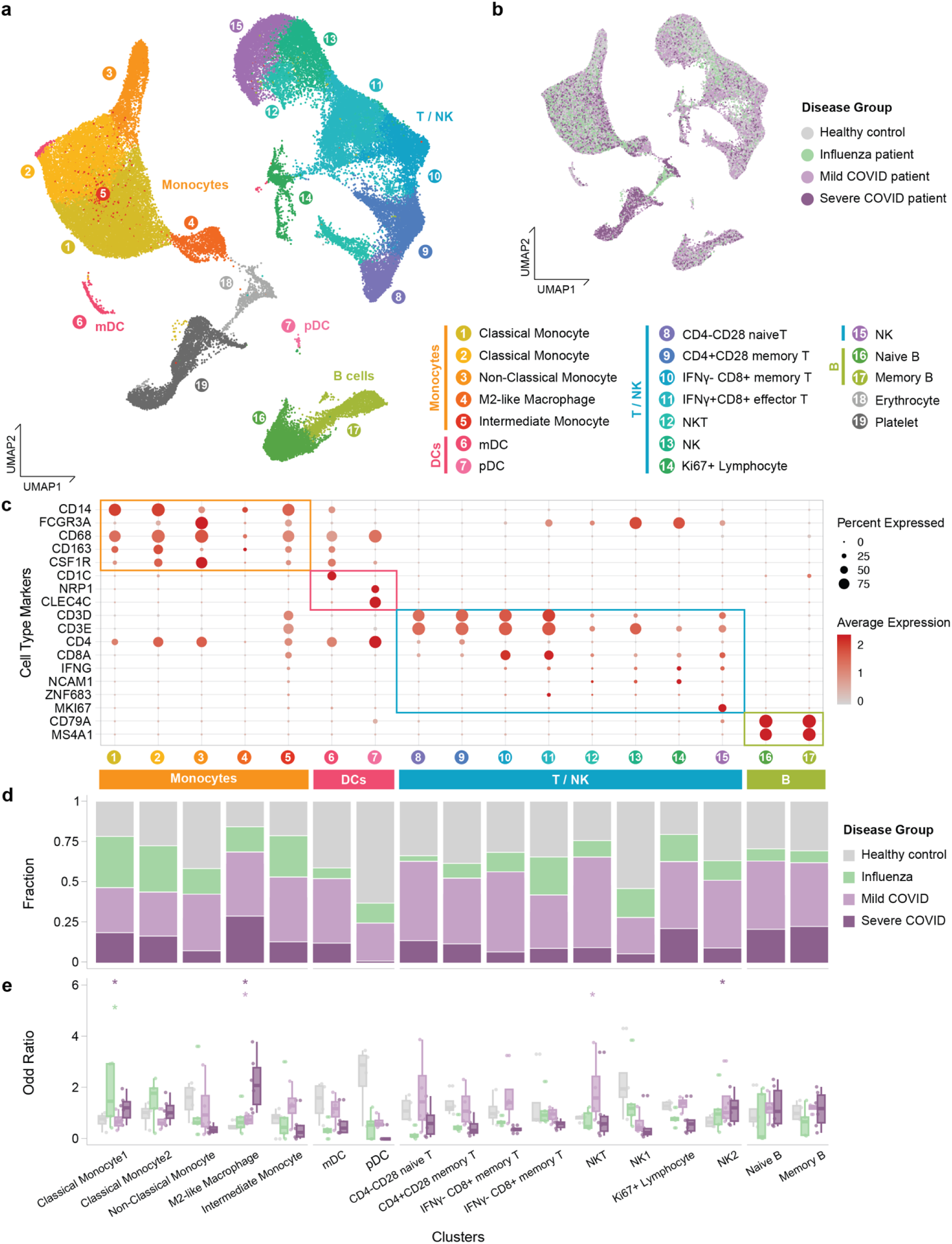
Overview of cell types in PBMC single-cell transcriptome of influenza/COVID-19 patients or healthy controls from Lee dataset. Analysis of 62,290 PBMC single-cell transcriptomes from the Lee dataset. **a.** UMAP plot of all cells, colored and annotated by major cell type clusters **b**.UMAP plot colored by disease group. **c.** Dot plot showing the expression of canonical marker genes across the cell clusters. Dot size is proportional percentage of cells in each cluster expressing the marker gene, while the color gradient represents the average expression level. **d.** Stacked bar plot showing the fractional composition of each cell cluster by disease group. **e.** Box plots displaying the sample-wise odds ratio (OR) for cell enrichment in each cluster, comparing disease groups to healthy controls. Asterisks indicate clusters with notable enrichment trends (T-test, p-value < 0.2), such as the expansion of Classical Monocyte1 in influenza patients.

To assign biological identities to these clusters, we employed a dual-strategy approach. First, we examined the expression of canonical marker genes, which allowed for the broad categorization of clusters into major immune lineages such as T cells (*CD3D*), B cells (*MS4A1*), NK cells (*NCAM1*), and myeloid cells (*CD14, FCGR3A*) (**Figure 2c**). This was supplemented with reference-based cell type annotation methods (SingleR, see Methods) to further refine these identities into more granular subtypes (**Table 1**, **Supplementary Figure 1**). This approach successfully identified populations of immune cells including T-cells, NK cells, B-cells, monocytes, myeloid/conventional dendritic cells (mDC), plasmacytoid dendritic cells (pDC), platelets and red blood cells/erythrocytes.

To assess how the proportions of these cell types shifted with disease status, a quantitative compositional analysis was then performed (**Figure 2d**). We evaluated the enrichment of each cell cluster across samples to assess this heterogeneity. Sample-wise odds ratios (OR) were compared between disease group (influenza or COVID-19) and healthy controls using T-test to examine the robust enrichments of each cell cluster in a diseased status (**Figure 2e**). For influenza infected patients, we observed moderate expansion of Classical Monocyte1 with consistently higher OR from influenza infected samples than healthy controls (2.45-fold increase, p = 0.122). For severe COVID-19 patients, we also observed expansions of Classical Monocyte1 (cluster 1; T-test p-value=1.11E-1, 1.68-fold increase), M2-like macrophages (cluster 4; T-test p-value=1.13E-2, 4.29-fold increase) and Ki67+ lymphocytes (cluster 14; T-test p-value=1.00E-1, 1.73-fold increase). For mild COVID-19 patients, we observed expansions of M2-like macrophages (cluster 4; T-test p-value=8.94E-2, 1.51-fold increase) and NKT cells (cluster 12; T-test p-value=1.92E-2, 2.38-fold increase). These results indicate that Classical Monocyte1 expansion is observed in both influenza and severe COVID-19, whereas M2-like macrophages show an enrichment specifically COVID-19 patients with more mild disease.

We conducted an integrative single-cell network analysis on the preprocessed data to highlight its underlying regulatory mechanisms and to investigate novel therapeutic targets for these infectious diseases. Briefly, differentially expressed genes (DEGs) were first identified for each cell cluster. Then, followed by the construction of gene co-expression networks to systematically characterize cell cluster-specific gene network models and co-expressed modules. These modules were integrated with the corresponding DEG signatures, which were mapped onto the respective network models. Enrichment analyses were performed to identify dysregulated cell type-specific subnetworks, and to determine potential upstream regulators by assessing DEG signatures within highly connected network hubs (i.e. key drivers)

### Cell type-specific network analysis revealed that the Monocyte and mDC populations are responsible for driving pro-inflammatory pathways

To pinpoint the most transcriptionally active cell populations, cell cluster-specific differential gene expression analysis (DEA) was performed. We compared cellular responses in samples from donors experiencing disease (influenza or COVID-19) against healthy control cells, and severe against mild COVID-19. This analysis confirmed that the most profound transcriptional perturbations in response to viral infection occurred within myeloid compartment (***Supplementary Figure 2***). Monocyte and mDC populations exhibited a large number of DEGs, indicating a high degree of activation and functional change. In contrast, lymphoid populations, such as T and B cells, displayed a markedly muted transcriptional response with far fewer DEGs.

Across Classical Monocytes and Non-Classical Monocyte, mainly up-regulated responses were observed during influenza and COVID-19 and responses between mild vs. severe COVID-19 were more balanced. These Classical and Non-Classical Monocytes showed similar expression patterns within Influenza and SARS-CoV-2 infection scenarios (**Figure 3**a). They shared pro-inflammatory and innate immune response signatures prominently. During influenza infection, commonly up-regulated genes include *ADM (Adrenomedullin)*^15^, *SLC39A8 (Solute Carrier Family 39 Member 8)*^16^, and *TLR2* (*Toll Like Receptor 2*)^17^, indicating active inflammatory signaling. On the other hand, *ITGA4 (Integrin Subunit Alpha 4)* was consistently downregulated which may have led to an increased recruitment of peripheral monocytes in influenza infection^18^. In COVID-19 case, Classical and Non-Classical Monocytes demonstrated an up-regulation of inflammatory modulators *IL-10 (Interleukin 10)*, *CD82 (CD82 Molecule)*, *TNFAIP6 (TNF Alpha Induced Protein 6)*, *CCRL2 (C-C Motif Chemokine Receptor Like 2)*, *CD55 (CD55 Molecule)* reflecting a similar response pattern also observed in influenza infection case. Additionally, chemokines such as *CXCL8 (C-X-C Motif Chemokine Ligand 8)* were commonly upregulated, which indicates strong innate immune activation. M2-like Macrophage and Intermediate Monocyte showed more scenario-specific response than Classical and Non-Classical monocytes. M2-like Macrophage exhibited significant overlap between genes up-regulated in COVID-19 but down-regulated in influenza infection. Intermediate monocyte uniquely responded to COVID-19 cases not evident in influenza or severe COVID-19 scenarios, highlighting a COVID-19 specific response.

**Figure 3.**
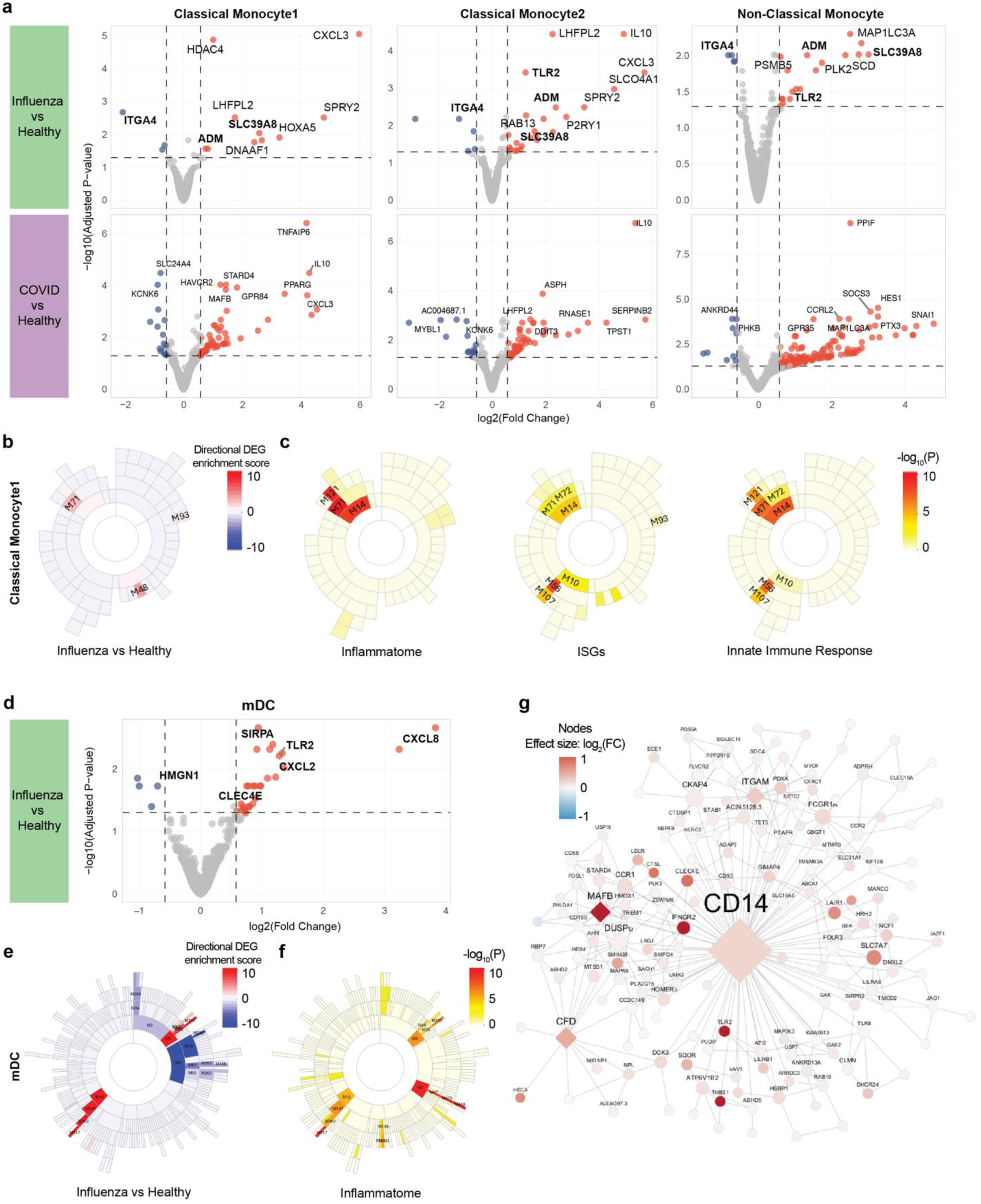
Differential expression and network analyses highlight pro-inflammatory responses in monocytes and mDCs. **a.** Volcano plots displaying differentially expressed genes (DEGs) in Classical Monocyte 1, Classical Monocyte 2, and Non-Classical Monocyte clusters. Comparisons are for influenza vs. healthy (top) and COVID-19 vs. healthy (bottom). Red and blue dots indicate significantly up- and downregulated genes (FDR < 0.05, |Fold Change (FC)| > 1.2), respectively. **b.** Sunburst plots showing directional DEG enrichment for co-expression modules from Classical Monocyte 1 network. The color reflects a directional enrichment where red indicates enrichment of upregulated DEGs (positive) and blue with downregulated DEGs(negative). The color intensity scales with statistical significance, highlighting modules with this score (see Methods). **c.** Sunburst plots showing functional enrichment for the modules in **b** in key immune pathways, including inflammasome, interferon-stimulated gene (ISG), and innate immune response pathway genes. Color intensity corresponds to the enrichment p-value. **d.** Volcano plot of DEGs in mDC cluster (influenza vs. healthy), highlighting the significant upregulation of innate immune genes including CLEC4E. **e.** Corresponding sunburst plot for mDC co-expression modules, showing directional DEG enrichment and **f.** functional enrichment in the inflammasome pathway. **g.** Co-expression module M50 from mDC network, which was found to be significantly enriched with upregulated DEGs and inflammasome related genes. Nodes represent genes and edges indicates significant co-expression relationships. Hub genes are denoted by a diamond shape. The size of each node is proportional to its degree and color represents the log2(FC) in expression for influenza vs healthy.

Having identified monocytes and mDCs as the key responding cell types by number of DEGs, we next sought to move beyond individual gene changes to understand the underlying regulatory architecture governing their responses. To achieve this, we constructed cell clusters-specific gene co-expression networks (see Methods), which coordinated expression for genes and can reveal functional modules and their key drivers. In Monocyte clusters, network analysis identified numerous distinct co-expression modules. We then integrated our DEG results with these networks using enrichment analysis to assess which modules were most active during infection (**Figure 3**b). This identified that modules significantly and robustly upregulated in influenza patients were functionally enriched for critical innate immune pathways, including inflammasome activation and interferon-stimulated gene (ISG) signaling. This confirmed that the transcriptional signature in monocytes reflected a coordinated pro-inflammatory program.

A parallel analysis in mDC population revealed an even more distinct and potent response to influenza. The mDC DEG signatures were particularly pronounced during influenza infection and severe COVID-19, but not between COVID-19 and control. Influenza infected mDCs was characterized by the strong upregulation of innate immune genes, including *CLEC4E*, alongside proinflammatory chemokines such as *CXCL8*, and other pattern recognition receptor like *TLR2* (***Figure 3***d). Downregulated genes included regulatory elements like *HMGN1* and *SNHG32*, potentially modulating the severity of the immune response. In contrast, the severely COVID-19 mDCs involved upregulation of neutrophil degranulation and innate immune-related genes, such as S100 family genes, *THBS1, CLU,* alongside downregulated antiviral immune markers (*FCER1A, CD1C, PLD4*) and MHC-II genes (*HLA-DMA, HLA-DRA*), indicating immune dysregulation. We then performed the same co-expression network generation and directional network enrichment analysis on mDC cluster. The results showed a dominant upregulation of modules functionally annotated with the inflammasome gene set (***Figure 3***e, f). To identify the upstream regulators modulating this response, we focused on the most significantly upregulated and functionally relevant module, M50. This module was not only highly enriched with inflammasome-related genes, but network topology analysis also found some genes including *CLEC4E* and CD14 were positioned as a central hub gene, characterized by a high degree of connectivity to other genes in the module (***Figure 5g***).

Integrative network analysis highlights conserved and infection-specific immune response. Influenza infection was characterized by IFN-driven innate immune activation, while COVID-19 involved pro-inflammatory and neutrophil-driven innate responses.^19^ Notably, monocytes and mDC clusters consistently displayed enriched pro-inflammatory signatures across both infections. Given that the pro-inflammatory signature in mDC was particularly potent and distinct in influenza, characterized by a strong innate immunity activation and pro-inflammation, we focused our subsequent analysis on influenza infected cohort to identify the core regulatory drivers of this pathogenic response.

### Multi-layered screening and in silico validation confirm CLEC4E’s robust pro-inflammatory signature during influenza infection

To systematically identify novel and clinically relevant therapeutic targets that driving pro-inflammatory response within monocyte and mDC populations, we applied an integrated multi-layered analysis. This approach was designed to integrate and score multiple independent lines of evidence to increase the confidence in our predictions. Each candidate gene was ranked based on composite score derived from: its significance as single-cell cell-type and condition-specific DEGs, its status as a key driver (KD) in our network analysis, its significance as pseudo-temporal DEGs, its correlation with clinical disease traits from publicly available bulk RNA-seq data, and novelty score based on its appearance in existing scientific publication related to influenza.

In brief, we identified potential key drivers (KDs) from the previously generated cluster specific network. Then, we integrated these single-cell derived signatures (DEGs and KDs) with bulk transcriptomic dataset from infected patients, emphasizing the correlation of candidate gene expression with clinical traits and disease severity. We also screened the identified candidate targets against existing literature, assessing the frequency of their appearance in publication titles and abstracts together with terms “influenza” to ensure novelty of the targets. Targets were ranked by a composite scoring system reflecting their significance as DEGs in scRNA-seq data, concordance with bulk data, and prominence as key drivers within cell-specific co-expression networks. We prioritized highly ranked, upregulated key drivers that exhibited strong specificity for monocyte and mDC subsets, minimal expression in other immune populations, and clear association with pseudotime trajectory-driven lineage differentiation during infection. Through this unbiased screening we could get top 10 promising candidate targets including *CLEC4E, VCAN, UPP1, CLEC10A, AQP9, NAMPT, SLC7A7, SDCBP, ALDH2, ACSL1* (**Figure 4**a).

**Figure 4.**
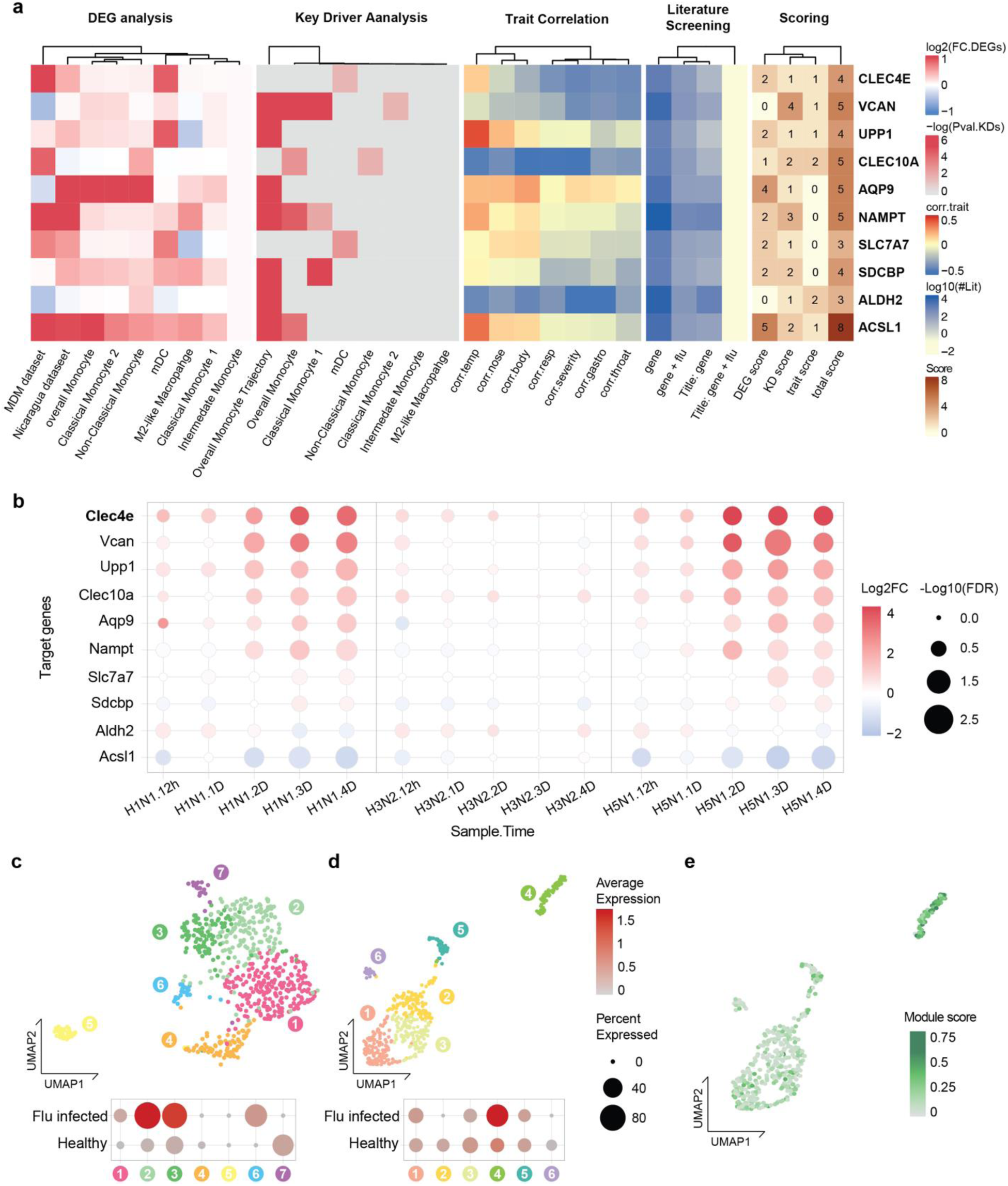
Prioritization of novel key driver through multi-layered analysis and in silico validation identifies CLEC4E as a top target. **a.** Heatmap summarizing the scoring of top candidate genes. Rows represent candidate genes, while columns represent different lines of evidence, including DEG and KD analysis, correlation with clinical traits, and literature novelty scores (see Methods). b. *in silico* validation using public bulk RNA-seq data from the lungs of mice infected with influenza strains of varying severity. Dot color represents log2FC and size indicates statistical significancy (FDR), showing robust upregulation of CLEC4E. **c.** In-depth analysis of mDC population in Lee dataset. UMAP of re-clustered mDC population (top). Dot plots showing specific overexpression of CLEC4E in subcluster 2 and 3 during influenza infection (color: average expression; size: percent of expressing cells). **d.** Validation of CLEC4E overexpression in independent scRNA-seq data generated from human influenza infected and healthy control PBMC samples. UMAP of re-clustered mDC population (top) and corresponding dot plot confirming CLEC4E overexpression in a specific mDC subcluster 4 during infection (bottom). **e.** UMAP of independent dataset (from **d**) with projected module score. The score, derived from marker genes of CLEC4E-overexpressed subcluster (2and 3) from Lee dataset, is high in the same patterned subcluster. This plot confirmed that the conserved transcriptional identity of this mDC subpopulation across datasets.

We then conducted a multi-step *in silico* validation to confirm the robustness of this finding. First, we interrogated public bulk RNA-seq data from the lung of mice infected with influenza strains of varying severity (ranked by increasing severity: H3N2, H1N1, H5N1). *Clec4e, Vcan, Upp1, Clec10a, Aqp9,* and *Nampt* exhibited an upregulated expression pattern in infected cases, consistent with the trends observed in the Lee dataset. Among them, *Clec4e* showed pronounced upregulation in response to severe strains (H1N1 day4: 3.41-log2fold increase, -log10(FDR)=1.2, H5N1 day4: 4.14-log2fold increase, - log_10_(FDR)=1.16), with expression levels increasing progressively over time post-infection. In contrast, *Acsl1* was downregulated in infected conditions within this dataset, suggesting that its expression may differ between blood and lung tissue or between species^20^ (**Figure 4**b).

Next, we validated these findings using publicly available single-cell RNA-seq data from influenza infected human PBMC samples. We began by in-depth re-clustering of mDC population from original Lee dataset. This higher-resolution analysis identified seven distinct mDC subclusters. Notably, *CLEC4E* was not uniformly expressed but was specifically overexpressed in two related subclusters (cluster 2 and 3) - and there only in samples from influenza patient, not in healthy controls (***Figure 4***c). To ensure this was not an artifact of a single dataset, we sought to replicate this finding in completely independent human influenza PBMC dataset^14^. We performed same mDC sub-clustering in this independent dataset and observed the identical results, specific mDC subcluster (cluster 4 in this dataset) also showed influenza infection-specific *CLEC4E* upregulation (***Figure 4***d). We then created a unique gene signature score based on the marker genes of the *CLCE4E*-high subclusters from Lee dataset and projected it onto the independent dataset, to prove that *CLEC4E*-high cells from separate studies were indeed the same cell population. The score was specifically enriched in the very same subcluster that we had identified as being *CLEC4E*-high and influenza-specific (***Figure 4***e).

### Pharmacological inhibition of CLEC4E ameliorates disease severity *in vivo*

Based on the strong *in silico* evidence supporting *CLEC4E*’s robustness, we lastly evaluated the *CLEC4E*’s potential as a therapeutic target *in vivo*. We utilized a murine model of influenza A (H1N1) infection and treated mice with the piceatannol, a known pharmacological inhibitor of Syk kinase which is a critical downstream component of *CLEC4E* signaling (***Figure 5***a). As hypothesized, mice treated with piceatannol exhibited significantly ameliorated disease severity. They experienced substantially less weight loss compared to untreated infected mice over the course of infection (***Figure 5***b). Consistent with the improved clinical outcome, piceatannol treatment also resulted in a significant reduction in viral titers in the lungs at day 5 post-infection (***Figure 5***c).

**Figure 5.**
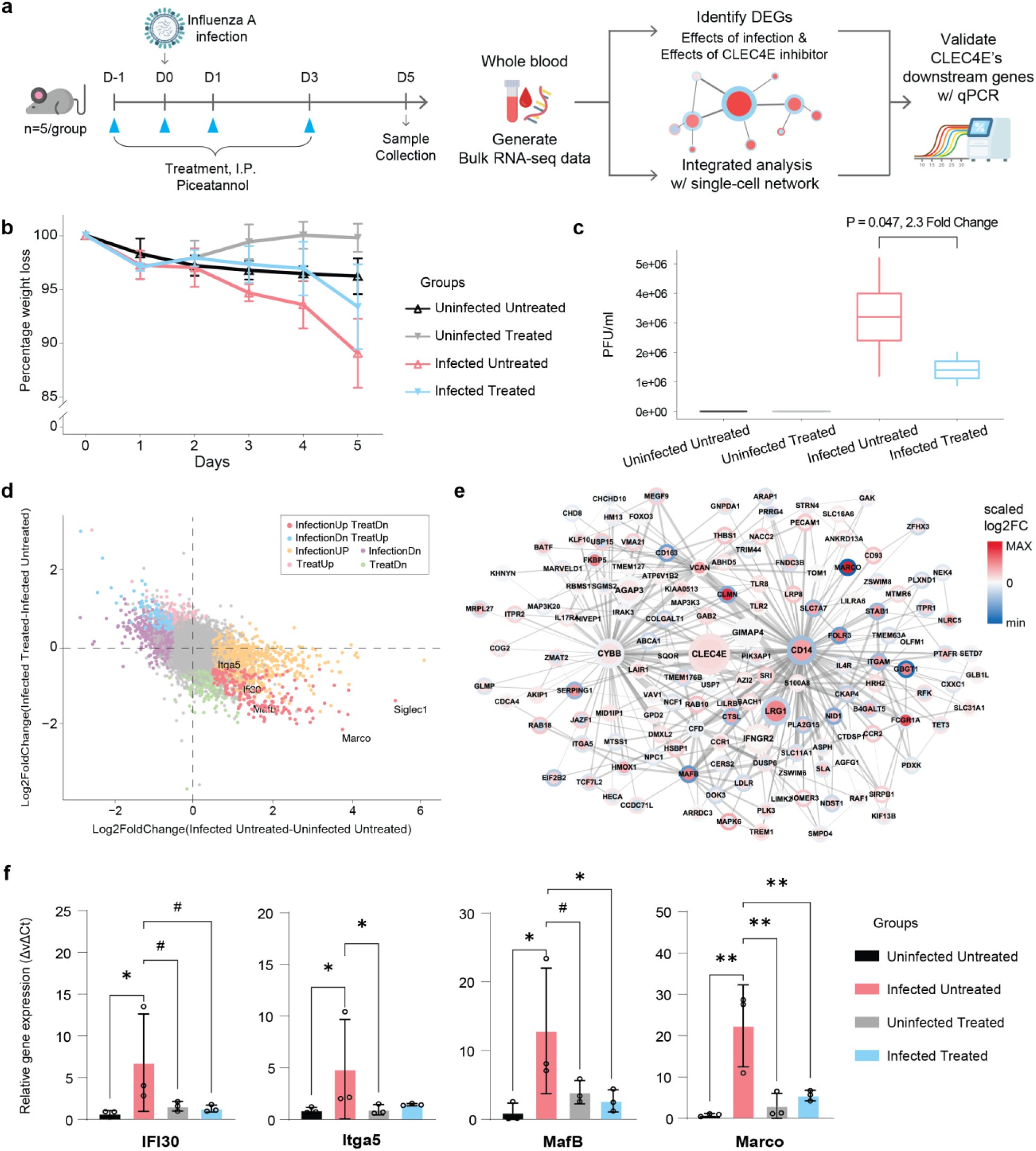
Targeting the inflammatory modulator CLEC4E alleviates disease severity *in vivo*. **a**. Schematic overview of the *in vivo* validation experiment. Mice were infected with influenza A virus and subsequently treated with Piceatannol (CLEC4E inhibitor) or a mock drug (PBS). Whole blood was collected on day 6 for bulk RNA-seq and qPCR. **b**. Daily body weight changes in four experimental groups: Uninfected Untreated(n=5), Uninfected Treated(n=5), Infected Untreated(n=5) and Infected Treated(n=4). **c**. The box plot shows viral titers (PFU/ml) in lung of each group’s mice at day 5. The median (center line), interquartile range (box), and min/max values (whiskers) (n per group are the same as panel b) **d**. Fold change–fold change plot shows DEGs. The x-axis represents expression changes due to infection, and the y-axis represents expression changes due to Piceatannol treatment during infection. **e**. **CLEC4E**-centered co-expression network generated from Lee dataset mDC cluster. Node color indicates the scaled log_2_fold change between infected and control groups (inner circle) or Piceatannol-treated and untreated group in infected mice. **f**. qPCR validation of selected CLEC4E downstream DEGs. Relative gene expression was calculated with 18S as the housekeeping gene. Data are presented as mean ± s.e.m. (n=3 per group). Statistical significance was determined using t-test for panel C and one-way ANOVA with Tukey’s post hoc test for f. # p < 0.07, * p < 0.05, ** p < 0.001.

To verify that piceatannol’s protective effects result from inhibition of CLEC4E-mediated pro-inflammatory programs, we collected blood from all four mouse groups after treatment and performed bulk RNA-seq. We quantified infection-driven and treatment-driven transcriptional changes separately, this fold change-fold change analysis revealed a set of genes whose upregulation during infection was specifically reserved by piceatannol treatment (***Figure 5***d). When we mapped these reversed DEGs back onto our Lee mDC co-expression network, we found that they were significantly enriched within *CLEC4E*’s immediate network neighborhood (***Figure 5***e). This analysis identified a statistically significant set of 20 genes that are infection-upregulated, treatment-downregulated, and reside within *CLEC4E*’s network neighborhood, including *IFI30*, *ITGA5*, *MAFB*, and *MARCO*. qPCR validation of selected genes confirmed their infection-induced upregulation and subsequent suppression by piceatannol treatment (***Figure 5***f). Collectively, our validations provide strong *in vivo* evidence that *CLEC4E* is a key functional driver of influenza pathogenesis and a promising therapeutic target for mitigating severe disease.

## Discussion

In this study, we first dissected the cellular response reflected in the blood during COVID-19 and influenza infection by analyzing single-cell transcriptomes of PBMCs from infected patients and healthy controls. This showed monocytes and mDCs as the primary cellular drivers of pro-inflammatory responses in blood. To identify the molecular regulators within these cell types, we applied cell type-specific co-expression network analysis. By integrating co-expression network modules with differentially expressed signatures and novelty scores, we highlighted *CLEC4E*, C-type lectin receptor, as a top novel pro-inflammatory driver in mDCs during influenza infection. We then performed a multi-step validation. *in silico* validation confirmed that *CLEC4E* was selectively upregulated in mDC subpopulation across independent datasets. Then, we tested its functional importance in vivo using a murine influenza model. We found that pharmacological inhibition of *CLEC4E* led to ease in disease severity via treated mice showed less weight loss and significantly lower lung viral load.

Our findings place *CLEC4E*, coding for the protein known as Mincle, as a key mediator of immunopathology in a new context. This C-type lectin is well-characterized as a pattern recognition receptor for exogenous pathogen-associated molecular patterns (PAMPs) which primarily from fungi and bacteria, as well as for endogenous damage-associated molecular patterns (DAMPs) which released from necrotic host cells. It typically drives inflammation through the FcRγ-Syk-CARD9 signaling pathway. While its role in sensing direct viral PAMPs is less clear, its function as a DAMP sensor suggests a mechanistic connection to influenza induced immune pathology, which is linked by extensive tissue damage and cell death. We propose a primary model where *CLEC4E* on mDCs senses DAMPs from virus-damaged lung tissue, triggering a pathological inflammatory cascade. This raises the question of how this localized response in lung manifests in PBMC analyzed in our study. One possibility is that DAMP from severe lung injury leaks into the bloodstream, where they directly activate circulating or bone marrow-derived mDCs via *CLEC4E*. Alternatively, systemic pro-inflammatory cytokines originating from the lung could prime mDC precursors, leading to an upregulation of *CLEC4E* and hyper-responsive state in the periphery. While the exact mechanism needs further investigation, our study provided compelling evidence that its sensing mechanism is also a pivotal component of the pathological response to a viral pathogen. Crucially, we demonstrated that *CLEC4E* is not merely a marker of inflammation but potentially can act as a key functional driver of disease.

Beyond the biological discovery of *CLEC4E*, our study provides a significant methodological contribution to the field of infectious disease research. Though some network analyses have been performed on bulk transcriptomic data from influenza patients, the application of co-expression network analysis at single-cell resolution to discover cell-type specific regulatory systems has remained underexplored. The power of our approach is underscored by the fact that we identified *CLEC4E* as a novel key driver by re-analyzing a public dataset, a critical regulator that was not highlighted in the original study which relied more on conventional differential expression analysis. This was not achieved by a single method, but by an integrated framework that synergized cell type-specific DEGs, network construction, key driver analysis, pseudo-temporal analysis and literature screening. We argue that this multi-layered framework serves as a robust and effective blueprint for mining novel therapeutic targets from future single-cell datasets not only for influenza study but for other complex diseases as well.

While our study provided strong functional validation in murine model, we acknowledge limitations that highlight directions for future research. For potential clinical application, the development of highly specific *CLEC4E* antagonists is also essential. Even though piceatannol proved effective, finding other therapeutic agents with improved specificity would be required to minimize potential off-target effects. Finally, it is important to determine if this CLEC4E-mediated immunopathology contributes to other severe viral infections. Our cross-disease network analysis provides preliminary evidence that this pathway may be relevant beyond influenza; we observed a significant enrichment of the *CLEC4E* in mDCs from COVID-19 patient as well (*Supplementary* Figure 5). Therefore, confirming the role of *CLEC4E* in COVID-19 pathogenesis would substantially broaden the therapeutic applicability of targeting this pathway.

In conclusion, our work identifies *CLEC4E* as a critical node in the pathological inflammatory network of influenza and validates it as a high-priority target for host-directed therapy. Modulating the activity of *CLEC4E* in mDCs represents a promising strategy to mitigate the excessive inflammation that drives severe disease, without globally suppressing the immune system. Ultimately, this study underscores the power of systems immunology to translate complex high-dimensional data into clinically relevant biological insights and actionable therapeutic strategies.

## Methods

### Lee Dataset Processing

The dataset from GSE149689, quantified and aggregated using the CellRanger pipeline, was downloaded for this project and reprocessed from scratch. Data was processed using the Seurat package (ver. 3.9.9)^21^. Quality control measures were applied to select high-quality cells based on mitochondrial gene expression rate (<20%) and the number of expressed genes (200-10,000). After stringent quality control steps, a total of 62,290 cells were retained. The count matrix was log-normalized with a scale factor of 10,000 using the “NormalizeData()” function. 2000 of the most variable genes were identified for each sample using “FindVariableFeatures()” with Variance Stabilizing Transformation (VST). Sample-wise data were integrated into a combined dataset through Seurat’s integration workflow^21^. “FindIntegrationAnchors()” was employed within the first 20 principal components to search robust anchor cells across different samples, followed by data merging with “IntegrateData()”. A shared nearest neighbor network (SNN) was constructed with “FindNeighbors()”. Clustering was performed using Louvain clustering at a resolution of 0.5 by “FindClusters()” and 19 clusters were identified. The Lee dataset^13^ divided patients into 5 disease status; Healthy control, Influenza patient, Asymptomatic case of COVID-19 patient, mild COVID-19 patient and severe COVID-19 patient. However, we excluded asymptomatic COVID-19 patient samples in our downstream analysis after cell type identification to minimize potential confounding effects from their ambiguous clinical presentations, thereby ensuring a clearer comparison of the immune profiles between patients with clearly defined symptomatic statuses.

### Cell type annotation

Clusters were annotated into specific cell types through a two-step approach to achieve a more precise result. First, all major immune cell populations were identified by utilizing well-established cell type markers, including T cells (CD3+), B cells (CD19+), monocytes/macrophages (CD14+), platelets (PF4+), erythrocytes (hemoglobin), dendritic cells (NRP1+), and NK cells (NCAM1^high^) (**Error! Reference source not found.**a,c).

Second, to get a more refined version of cell subtypes, SingleR^22^ was used for inferring cell types based on comparison between the Lee dataset and pre-annotated reference data (Supplementary Figure 1). Two different datasets were utilized as reference sets. Human Primary Cell Atlas (HPCA)^23^ collection is a microarray data of broader blood cell types. Monaco^24^ collection derived from PBMC of healthy individuals. It allows detection of detailed immune subsets. Using both reference datasets allowed the correct identification of cell types as reported in the original report from Lee dataset^13^. The overall cell type designation is summarized in Table 1.

**Table 1.** Summary of cell type assignments to the cell clusters in Figure 2.

| Major Cell Type | Cluster name | Subset name |
| --- | --- | --- |
| Monocytes (CD19+) | Mono.1 | Classical (CD14+CD16-) |
|  | Mono.2 | Classical (CD14+CD16-) |
|  | Mono.3 | Non-classical (CD14dimCD16+) |
|  | Mono.4 | M2-like macrophage (CD68+CD163+) |
|  | Mono.5 | Intermediate (CD14+CD16+) |
| Dendritic Cells (NRP1+) | pDC | Plasmacytoid dendritic cells |
|  | mDC | Myeloid dendritic cells |
| T/NK lymphocytes<br>(CD3+/NCAM1+) | T1 | CD4- CD28+ naïve T-cells |
|  | T2 | CD4+ CD28+ naïve/memory T-cells |
| | T3 | IFN-γ<sup>-/low</sup> CD8+ memory T-cells |
| | T4 | IFN-γ+ CD8+ effector T-cells |
|  | T5 | NKT |
|  | T6 | NK |
|  | Prolif. | Ki67+ lymphocytes |
|  | NK | NK-cells |
| B lymphocytes<br>(CD79+) | B1 | Naïve B-cells |
|  | B2 | Non-switched memory B-cells |
| Platelets (PF4+) | PL |  |
| Erythrocytes (HBA1/2+) | RBC |  |

### Cluster specific differential expression analysis (DEA)

DEA was performed for each cluster to compare influenza- and COVID-19-patient oriented cells with healthy control cells. To improve reproducibility and reduce false-positive rates, we employed a pseudobulk-based approach^25,26^ which aggregates cell-wise read counts for each individual to generate cluster-specific pseudobulk data. DEGs were then identified using DESeq2^27^ with the model g_i_ ∼ disease group + age + gender + ɛ_o_. Disease groups included influenza-infected, mild COVID-19, severe COVID-19, and healthy controls, with each infected group compared to healthy controls. DEGs were considered significant with an FDR adjusted p-value < 0.05 and a fold change > 1.2 or < 1/1.2.

### Key Driver Analysis (KDA)

Key Driver Analysis (KDA) identifies regulatory nodes (drivers) for a given gene set *G* within a directed gene network *N*. It constructs a subnetwork *NG*, comprising nodes within *h* layers of *G*, with *h* determining the inclusion range—from a minimal subgraph (*h* = 0) to the entire network (*h* = ∞). Two strategies are used:

- Dynamic Neighborhood Search (DNS): For each candidate gene in *N_G_*, KDA computes enrichment statistics *ES_h,g_* for its h-layer neighborhood *HLN_g,_h*, selecting the optimal *h*\* that maximizes enrichment. Nodes with significantly enriched *HLN*s are designated as candidate drivers. Those without parent nodes in *N_G_* are termed global drivers. A secondary “outlier detection” step evaluates *HLN* enrichment against the full network *N*, rescuing key regulators missed in subnet-based tests.
- Static Neighborhood Search (SNS): Uses a fixed *h*-layer for all nodes. Nodes with *HLN* sizes significantly above the network mean *μ* + *σ*(*μ*), or without-degrees > *d* + 2*σ*(*d*), are nominated as candidate and global drivers, respectively.

This approach enables the systematic prioritization of regulatory hubs within the context of gene expression signatures and biological networks.

### Co-Expression Network Analysis

For cell cluster-specific co-expression network analysis, we have further processed the data to account for technical variability across different individuals. Specifically, we rescaled the average cell-wise read depth per sample to match the minimum across all samples using “rescaleBatches()” function from batchelor R package (v1.17.2)^28^. Multiscale Embedded Gene Co-Expression Network Analysis (MEGENA)^29^ was performed to identify single-cell clusters and pseudobulk host modules of highly co-expressed genes in the infection scenarios studied. The MEGENA workflow comprises the following major steps: (a) Fast Planar Filtered Network construction, (b) Multiscale Clustering Analysis, and (c) Multiscale Hub Analysis. The total relevance of each module to infection was calculated by using the Product of Ranks method with the combined enrichment of the differentially expressed gene (DEG) signatures as implemented.

### Module Enrichment Analysis

We employed Fisher’s Exact Test (FET) for enrichment analysis of co-expression modules for signatures such as directional DEGs, the inflammatome^30,31^, interferon stimulated genes (ISGs^32^), and genes of the innate immune response^33^ . P-values were adjusted for multiple testing (FDR) and considered significant with FDR ≤ 0.05. In the figures p-values were converted by -log_10_(P). Separately assessed FET p-values for up- and down-regulated DEGs (directional DEG enrichment) were combined as such: -log_10_(P_up_) + log_10_(P_down_). The limitation of this approach is that in the rare event of modules enriched for both up- and down-regulated DEGs the combined enrichment eventually cancels out.

### Trajectory Analysis

Given the importance of pro-inflammatory roles of monocytes in flu infections^13^, we sought to study the pseudo-temporal landscapes in monocytes underlying influenza infection. We utilized slingshot^34^ to infer cell trajectories amongst the monocytes to study cellular dynamics driven by different infection status. As the trajectory analysis requires initiating the starting cells for the trajectories, we first performed subclustering on the monocytes using shared nearest neighbor approach in Seurat^21^ with 20 principal components at γ=1.2. This yielded 18 subclusters (Supplementary Figure 3A). Subcluster 5 yielded the largest portion of healthy control cells and was utilized as the initial clusters for the subsequent trajectory analysis (Supplementary Figure 3B). As results, slingshot yielded 9 lineages (Supplementary Figure 3A). Then, we identified differentially expressed genes (DEGs) along the cell trajectory by adopting TradeSeq, which models non-linear gene expressions patterns along trajectory in generalized additive model (GAM) framework^35^. We performed an association test using ‘associationTest()’, and comparison of start and end cells using startVsEndTest() using the first 10% cells as the start and last 20% as the end cells across all lineages. The global p-values summarized across all lineages were utilized for identifying significant genes. After correcting the global p-values by Benjamini-Hochberg false discovery rate for multiple testing^36^, we applied FDR adjusted p-value ≤ 0.05 and fold change ≥ 2 to identify significant genes per test.

### Target Selection

We followed a multi-step process to identify cluster-specific targets with respect to influenza and COVID-19 cases. For each selected cluster, we identified cell-type and infection specific signatures including **DEGs** and **KDs**. We further assessed bulk-based gene / **trait** correlations. We used a public dataset of an influenza infection cohort (GSE114588^37^) that further included self-reported severity information. We also searched for literature references for the target genes. We included the latter type of information to evaluate the novelty of the predicted targets. We counted occurrences of target names in **literature** references, in the text, in the title, separate as well as in combination with the studied diseases.

Tally: Targets were ranked by compound scores based on occurrences as significant DEGs in single-cell data, co-occurrence as significant DEG in the bulk datasets, and single-cell KD status. Targets were filtered by low citation in literature, in particular in manuscript titles together with an of the terms “influenza”, “COVID” or “SARS-CoV-2”.

To prepare for further downstream experimental validation by drug induced inhibition, we focused on upregulated targets in monocyte and myeloid dendritic cell clusters with weak or no expression in other cell types. Our main priority in target selection was upregulated KDs in these specific clusters. We also accepted targets as upregulated KDs in overall monocytes and corresponding pseudotime trajectory. Additional selection criteria were significant correlations to clinical traits after cell type deconvolution^38^ of patient-based bulk transcriptome data. As we were further interested in novel targets, we selected against targets that have already been studied in connection with influenza, thus were mentioned in Europe PubMed Central titles together with the term “influenza”. We accepted mentioning the target name in the title by itself, or in the abstract (with or without cooccurrence with the term “influenza”).

### *In silico* Validation

For bulk RNA-seq validation, data from GEO under accession number GSE114588^37^ was downloaded and processed. The data consists of 56 samples derived from 34 subjects with time-points of an initial, infected and final, recovered phase. The R-packages *edgeR* was used to process the expression profile and to call DEGs using an FDR ≤ 0.05 and an effect size of |FC| ≥ 1.2.

For single-cell RNA-seq validation, the raw dataset GSE243629 was obtained from GEO database and processed using Seurat package (ver.5.1.0)^39^. This dataset contains 11 human PBMC samples, 7 of influenza A virus-infected patients, and 4 of healthy controls. As a quality control step, only cells with less than 20% of mitochondrial gene expression rate and more than 200 unique expressed genes and only genes which were expressed in more than 5 cells were kept. A total of 102,818 cells and 48,526 genes were used for further analysis. The count matrix was normalized with “SCTransform()” function while regressing out mitochondrial gene percentage as an unwanted source of variation. The top 2,000 most variable features were identified for downstream analyses. “IntegrateLayers()” function was applied to the sample-wise dataset using CCAIntegration method to correct batch effect between samples.

### Experimental Validation Design

Thirty female C57BL/6 mice (n = 5 per group, age: 6-8 weeks) were obtained from Jackson laboratories and housed under specified pathogen-free (SPF) conditions with ad libitum access to food and water. The experimental protocol was approved by the Institutional Animal Care and Use Committee (IACUC) of the Icahn School of Medicine at Mount Sinai and adhered to the guidelines for the care and use of laboratory animals. Mice were randomly assigned to four experimental groups:

Group 1: PBS (intraperitoneal, IP) + influenza virus

Group 2: Piceatannol (0.125 mg IP) + influenza virus

Group 3: Piceatannol (0.125 mg IP) + mock infection

Group 4: PBS (IP) + mock infection

### Virus Challenge

Influenza A virus (H1N1 strain IVR-180) was produced on embryonated eggs and titrated on Madin-Daraby Canine Kidney (MDCK) cells by plaque assay. Virus stocks were prepared as previously described^40^. Mice in the infected groups were challenged with a sublethal dose (0.5 Lethal Dose 50) of the virus diluted in PBS via the intranasal route in a volume of 50μL. Mice in the mock groups received an equivalent volume of sterile PBS. After infection, animals were monitored daily for 5 days for clinical signs of disease, body weight loss, and survival.

### Drug Treatment

Piceatannol was obtained from Thermofisher (328452500). Piceatannol (0.125 mg in PBS) was administered intraperitoneally (IP) in a total volume of 500μL (1 day before infection, the day of infection, and 1- and 3-days post infection for piceatannol). Mice in the PBS control group received an equivalent volume of sterile PBS via the same route and frequency.

### Sample Collection and Virus Titration

On day 5 post-infection, animals were euthanized by CO₂ inhalation followed by cervical dislocation. Lungs were excised, snap-frozen in liquid nitrogen, and stored at -80°C for virus titration by plaque assay as described elsewhere^40^ . Blood was collected via cardiac puncture into EDTA-coated tubes for RNA sequencing, following protocols optimized for RNA isolation from whole blood (Ribopure, InVitrogen).

### Bulk RNA sequencing and data processing

Sample QC, library preparations, sequencing reactions, and bioinformatic analysis were conducted at GENEWIZ, LLC./Azenta US, Inc (South Plainfield, NJ, USA) as follows:

### Sample QC

Total RNA samples were quantified using Qubit 4.0 Fluorometer (Life Technologies, Carlsbad, CA, USA) and RNA integrity was checked with 4200 TapeStation (Agilent Technologies, Palo Alto, CA, USA).

### Library Preparation and Sequencing

rRNA and globin depletion sequencing library was prepared by using QIAGEN FastSelect rRNA HMR and Globin Kit (Qiagen, Hilden, Germany). RNA sequencing library preparation used NEBNext Ultra II RNA Library Preparation Kit for Illumina by following the manufacturer’s recommendations (NEB, Ipswich, MA, USA). Briefly, enriched RNAs are fragmented for 15 minutes at 94 °C. First strand and second strand cDNA are subsequently synthesized. cDNA fragments are end repaired and adenylated at 3’ends, and universal adapters are ligated to cDNA fragments, followed by index addition and library enrichment with limited cycle PCR. Sequencing libraries were validated using the Agilent Tapestation 4200 (Agilent Technologies, Palo Alto, CA, USA), and quantified using Qubit 4.0 Fluorometer (ThermoFisher Scientific, Waltham, MA, USA) as well as by quantitative PCR (KAPA Biosystems, Wilmington, MA, USA).

The sequencing libraries were multiplexed and clustered onto a flowcell on the Illumina NovaSeq instrument according to manufacturer’s instructions. The samples were sequenced using a 2x150bp Paired End (PE) configuration. Image analysis and base calling were conducted by the NovaSeq Control Software (NCS). Raw sequence data (.bcl files) generated from Illumina NovaSeq was converted into fastq files and de-multiplexed using Illumina bcl2fastq 2.20 software. One mis-match was allowed for index sequence identification. The resulting raw sequencing reads in FASTQ format were first assessed for quality using FastQC^41^ (v.0.11.9). Adapters and low-quality bases were trimmed using fastp^42^ (v.1.0.0). The clean reads were aligned to the mouse reference genome (GRCm39) and quantified using STAR-RSEM pipeline (STAR^43^ v.2.7.11b, RSEM^44^ v.1.3.3) to generate a gene-level read count matrix.

DEA performed using R package DEseq2^27^. The count matrix was imported and genes with read counts ≥ 10 in at least half of the samples were retained for downstream analysis. To specifically isolate the effects of infection and treatment, we conducted two separate analyses. To determine the effect of infection, we analyzed only the PBS-treated samples (infected and uninfected) using design formula ∼Infection. DEGs were identified by contrasting the infected group against the uninfected group. To determine the effect of treatment, we used only influenza infected samples (piceatannol treated and PBS treated) with model ∼Treat, contrasting the treated group against untreated group. For both analyses, genes with FDR < 0.05 and absolute log2(FC) > 1 were considered statistically significant.

### qPCR

One step qRTPCR were performed on 10ng of RNA isolated from whole blood of three mice with Power SYBR™ Green RNA-to-CT™ 1-Step Kit (Thermofisher) as described elsewhere: 35590097 using the following primers:

**Table 2.**
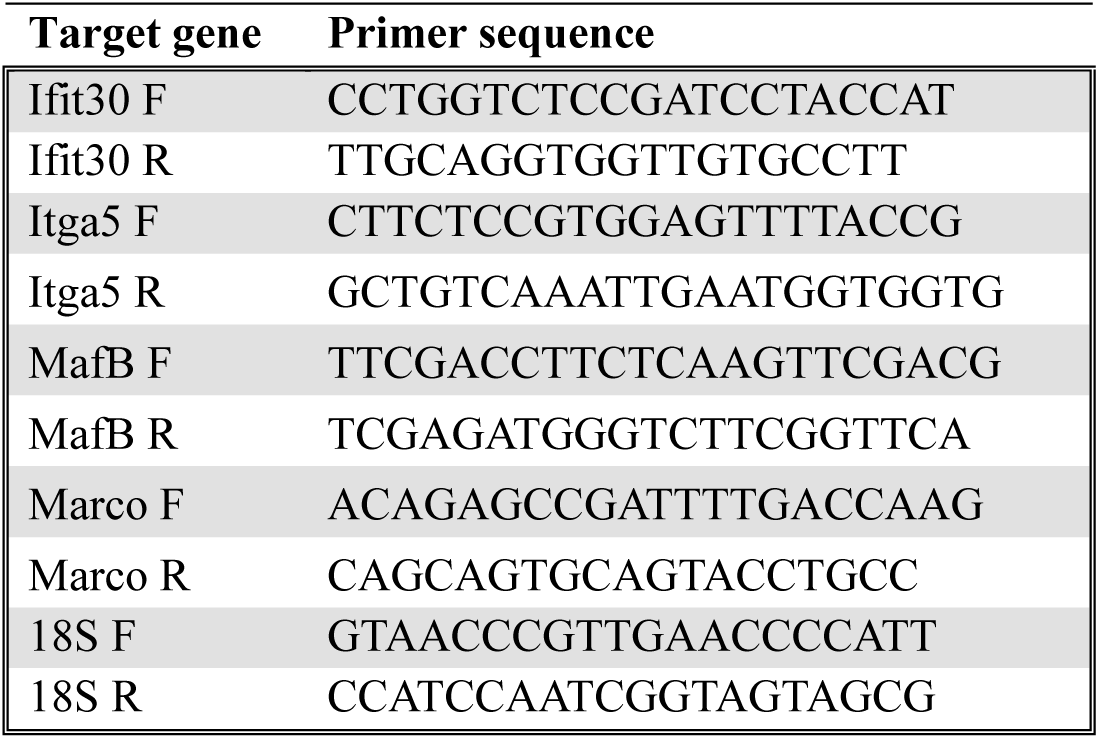
Primer sequence information.

18S was used as a housekeeping gene to normalize, and relative gene expression was calculated by 2^−ΔΔCt^ (GAPDH has been tested as housekeeping genes, but its expression pattern is not stable in the context of Influenza infection^45^)

## Acknowledgements

This work was supported by grants from the National Institute of Health (NIH) under award numbers R21AI149013, R01AI170112, U01AG088351.

This work was further supported in part through the Minerva computational and data resources and staff expertise provided by Scientific Computing and Data at the Icahn School of Medicine at Mount Sinai and supported by the Clinical and Translational Science Awards (CTSA) grant UL1TR004419 from the National Center for Advancing Translational Sciences.

## Author Contributions

Conceptualization: CVF, WMS; computational analysis: AM, CVF, SC, WMS; validation experiments: GL, GS, MS; writing-original draft: CVF, SC, WMS; writing-review and editing: AGS, CVF, GL, MS, SC, WMS; supervision: AGS, CF, MS, WMS; funding acquisition: AGS, CF, MS, WMS.

## Competing Interests

The M.S. laboratory has received unrelated funding support in sponsored research agreements from Phio Pharmaceuticals, 7Hills Pharma, ArgenX NV, Ziphius and Moderna.

---

**Supplementary S1.**
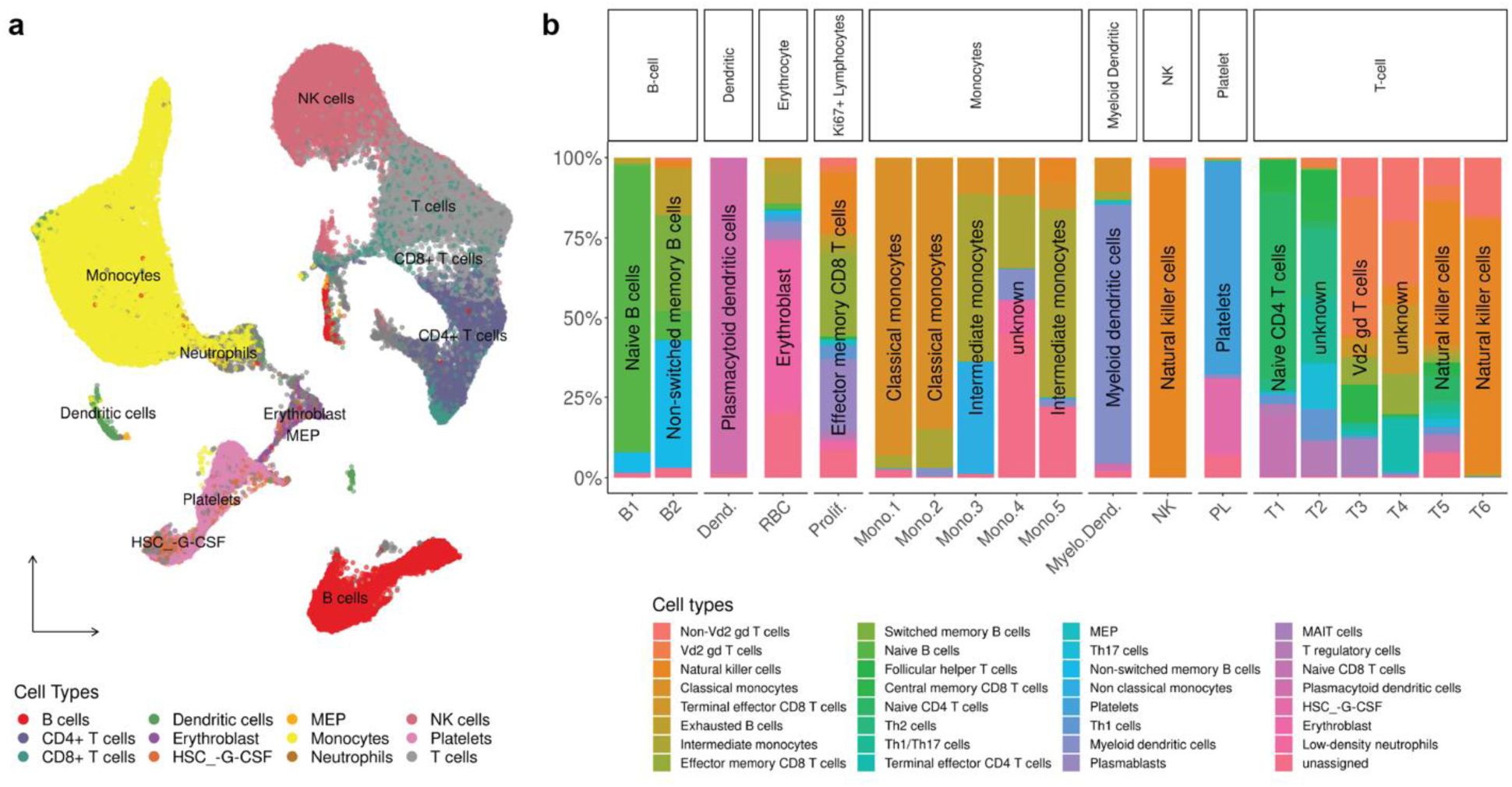
Summary of SingleR inferred cell types. A. UMAP plot of SingleR inferred results. Different colors depict different cell types shown on the legend below. B. Proportion bar plot of SingleR inferred cell subtypes in cell clusters. The most pronounced cell subtype with > 25% in each cluster is labeled. Cell clusters with no cell subtypes exceeding > 25% are labeled unknown.

**Supplementary S2.**
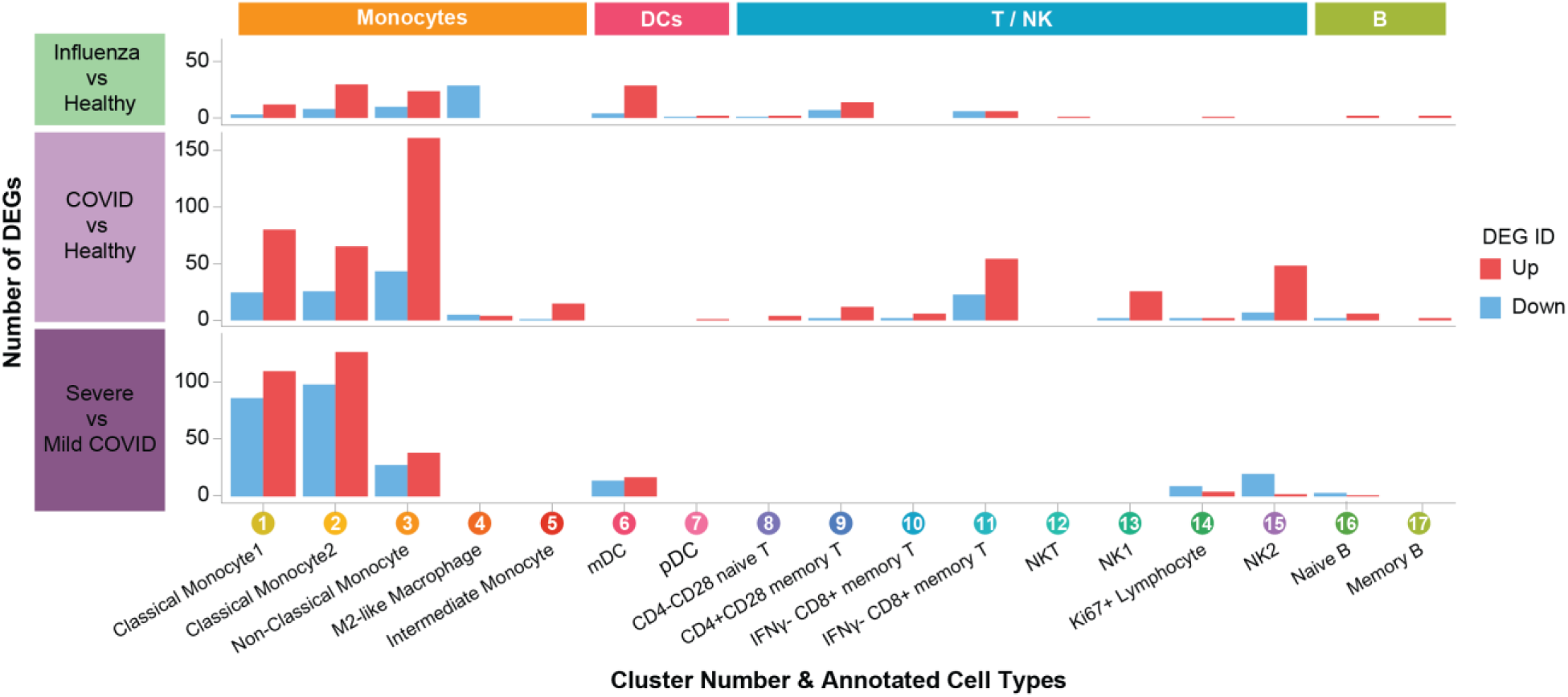
Number of significant differential expression genes (DEGs) with FDR < 0.05 and, FC > 1.2 or FC < 1/1.2 from individual cell clusters by different scenario (Influenza patient vs Healthy control, COVID patient vs Healthy control and Severe COVID vs Mild COVID patient)

**Supplementary S3.**
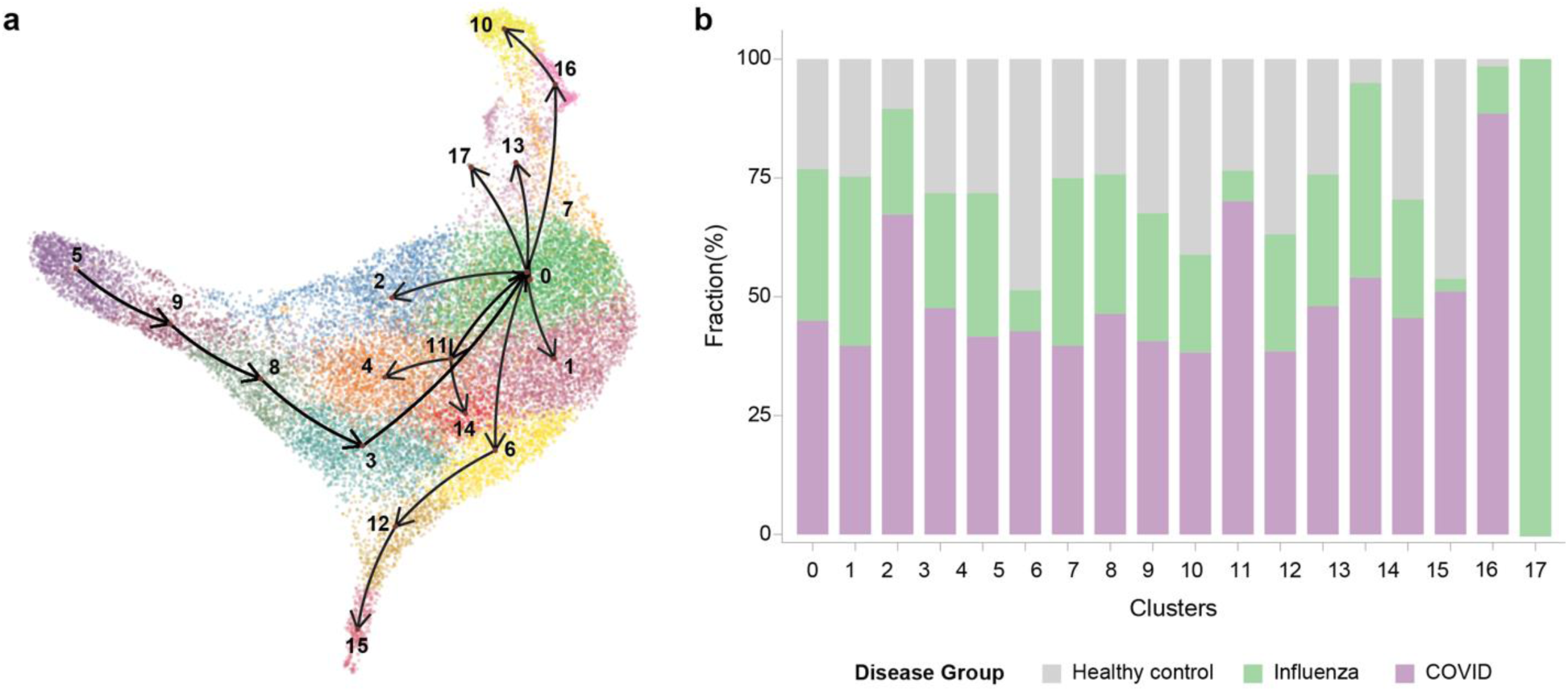
Monocyte trajectory analysis. A. UMAP plot of monocytes and its subclusters. The arrow shows trajectory progressions across 9 different lineages, inferred from setting subcluster 5 as the initial cells. **B. Proportions of different disease groups** in monocyte subclusters.

**Supplementary S4.**
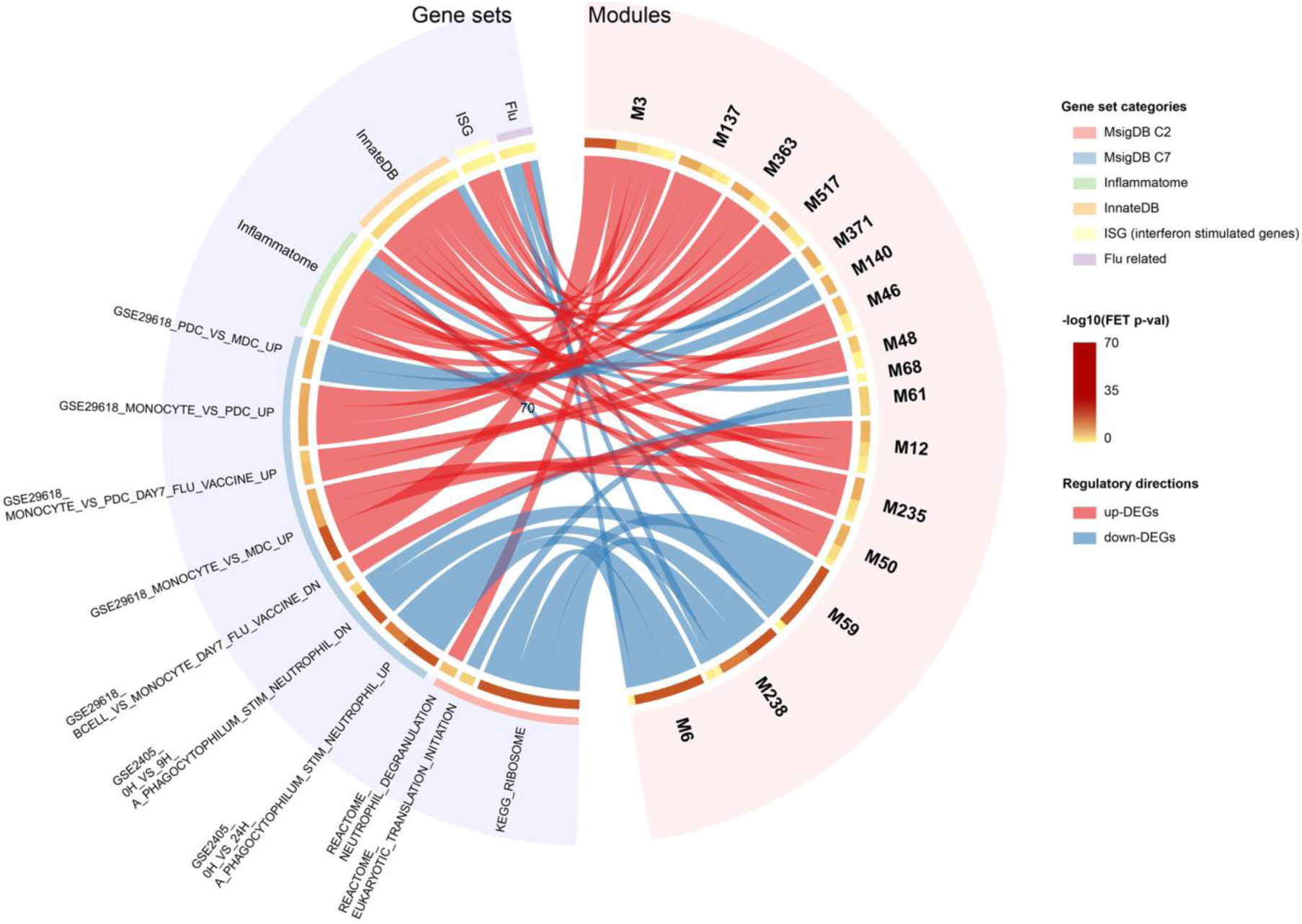
Directional functional enrichment of mDC co-expression modules. A circus plot visualizing the significant associations between co-expression modules identified in mDC cluster (right side, Modules) and major functional pathway categories (left side, “Gene sets”). The associations displayed were selected using a priority-base filtering approach to highlight the most relevant biological themes while maintaining clarify. Specifically, all significant associations for key immune related gene sets (Inflammatome, ISG, Innate immune and Flu) are included. To avoid visual clutter, only the top 20 most significant associations from the remaining pathway categories are displayed. Each link’s color indicates the overall regulatory direction of the module, determined by comparing enrichment for up-(red links) versus down(blue links)-regulated DEGs. The thickness of each link is proportional to the statistical significance (-log10(p-value) of the association. Modules on the right are ordered using hierarchical clustering to group those with similar functional profiles. The gene sets defining the immune-related pathways are provided in Suppl. Table 1. For a complete list of all significant module-pathway enrichments, see Suppl. Table 2

**Supplementary S5.**
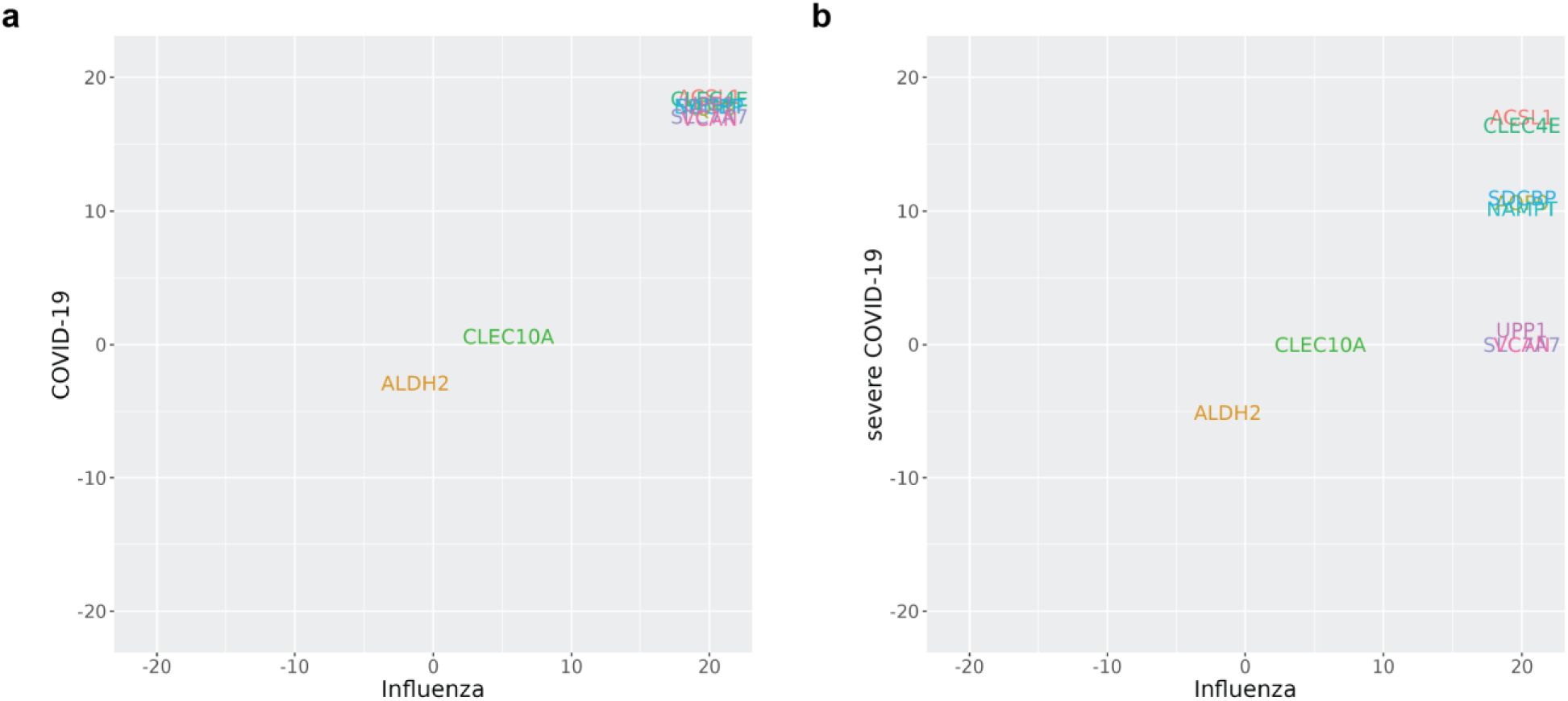
Disease-specific KDA of target KDs in mDC cluster. Negative log_10_ adjusted p-values of selected KDs after KDA using disease specific, directional DEGs as signatures are shown – -log_10_ transformed KDA p-values using up-regulated DEGs are positive, down-regulated DEGs are annotated with negative values. **a.** Target KDs during influenza (x-axis) and COVID-19 (y-axis) are shown. **b**. Target KDs during influenza (x-axis) and severe COVID-19 (y-axis) are displayed. Both influenza and COVID-19 DEGs are calculated against healthy control. All target KDs are required to be significantly upregulated (FC ≥ 1.5, FDR ≤ 0.05). For illustrative purposes KDA adjusted p-values are limited to ≥ 10^-20^ (-log_10_(P) ≤ 20). However, observed p-values of some of the KDs are as low as 10^-56^ (see Suppl. Table 3).

